# A Different Kind of Restraint Suitable for Molecular Dynamics Simulations

**DOI:** 10.1101/2022.08.27.505552

**Authors:** István Kolossváry, Woody Sherman

## Abstract

Conformational sampling of complex biomolecules is an emerging frontier in drug discovery. Indeed, advances in lab-based structural biology and related computational approaches like AlphaFold have made great strides in obtaining static protein structures. However, biology is in constant motion and many important biological processes rely on conformationally-driven events. Unrestrained molecular dynamics (MD) simulations require that the simulated time be comparable to the real time of the biological processes of interest, rendering pure MD impractical for many drug design projects, where conformationally-driven biological events can take microseconds to milliseconds or longer. An alternative approach is to accelerate the sampling of specific motions by applying restraints, guided by insights about the underlying biological process of interest. A plethora of restraints exist to limit the size of conformational search space, although each has drawbacks when simulating complex biological motions. In this work, we introduce a new kind of restraint for molecular dynamics simulations (MD) that is particularly well suited for complex conformationally-driven biological events, such as protein-ligand binding, allosteric modulations, conformational signalling, and membrane permeability. The new restraint, which relies on a barrier function (the scaled reciprocal function) is particularly beneficial to MD, where hard-wall restraints are needed with zero tolerance to restraint violation. We have implemented this restraint within a hybrid sampling framework that combines metadynamics and extended-Lagrangian adaptive biasing force (meta-eABF). We use two particular examples to demonstrate the value of this approach: (1) quantification of the approach of E3-loaded ubiquitin to a protein of interest as part of the Cullin ring ligase and (2) membrane permeability of heterobifunctional degrader molecules with a large degree of conformational flexibility. Future work will involve extension to additional systems and benchmarking of this approach compared with other methods.

## 1 Introduction

Molecular dynamics (MD) continues to grow in terms of utilization and impact in drug discovery programs [1–3]. In many cases, traditional unbiased MD simulations are insufficient to achieve the requisite level of sampling and accuracy within a time frame needed to make an impact in drug discovery projects. As such, using restraints in MD has become commonplace to improve convergence and sampling conformational states of interest. Restraining atomic positions or interatomic distances represents the most frequently used restraints, but many biologically relevant motions involve atomic transformations that cannot be treated effectively with such simple restraints and therefore require more sophisticated collective variables (CV) utilized in advanced sampling methods [4, 5], which often require more sophisticated restraints.

The dominant restraining algorithm used in MD today is a simple quadratic penalty function such as *F*_*c*_*(*x*−*x*_0_)^2^ where *x* is the variable to be restrained, *x*_0_ is its desired value, and *F*_*c*_ is a force constant. In this form the restraint penalty gets applied for all non-zero values (i.e. *x < x*_0_ or *x > x*_0_) and the restraint force is proportional to |*x* − *x*_0_|. Often we want to restrain a variable not to a particular value, but just want to make sure that the value does not exceed an upper limit and/or fall below a lower limit. This can be achieved by zeroing out the left hand side of the restraint parabola located at the upper limit, or zeroing out the right hand side of the restraint parabola placed at the lower limit. The combination of the two limits provides the ubiquitous “flat bottom” restraint that penalizes values under the lower limit and above the upper limit whereas no penalty is applied between the limits. Mathematically, “zeroing out” means multiplying the penalty function with a unit step function centered at the upper/lower limit, respectively.

Simple quadratic penalties as well as complex penalty functions involving higher powers, scaling, and offsets are used routinely in MD simulations. Note, however, that penalty based restraints allow the restrained value to violate the limits—by definition the penalty is only applied when the variables exceed the specified limits (if no offset is used). For many situations this approach is acceptable, but not in all cases—for example, if we want a restraint that cannot tolerate any violation. One well-known example is restraining covalent bonds involving H atoms to stay completely rigid in MD simulations to extend the time step beyond 1 fs without using multiple time step integration techniques [6, 7].

In the work presented here, we focus on upper/lower wall restraints in two types of advanced sampling simulations, one representing large-scale conformational transitions in proteins and the other estimating the potential of mean force (PMF) of large drug-like ligand molecules passing through a membrane. In both cases the wall must be impregnable. To that end, we utilize a class of so-called barrier functions. Barrier functions are designed to never let the restrained variable escape from its allowed domain by applying an infinite wall.

Barrier functions such as −*log*(*x*_0_ − *x*) or 1*/*(*x*_0_ − *x*) are staples of the so-called interior-point methods in constrained nonlinear optimization [8], where the attractive idea is that once the constrained value is inside the allowed domain, it can no longer escape because the barrier is infinite at the limit. Unfortunately, interior-point methods have two significant drawbacks that makes them difficult to use in optimization problems. One is that in general it is nontrivial to move the constrained variable inside the allowed domain in the first place. The main problem, though, is that general purpose line search methods can probe the exterior of the allowed domain where the barrier function either does not exists (logarithmic) or manifests a numerical “black hole” (reciprocal). The latter happens when the value of the escaped *x* gets arbitrarily close to the barrier wall from the outside where the barrier function goes to minus infinity. Modified line search methods do exist, but they only work efficiently with linearly constrained problems [9, 10]. Therefore, in most cases constrained nonlinear optimization problems are solved with penalty functions utilizing a series of successively increasing force constants [8], but that would be impractical in MD. The basic tenet of our thesis presented here is to demonstrate that reciprocal barrier methods do, in fact, work well in complex MD simulations.

Our barrier function of choice is the simple reciprocal *k/*(*x*_0_ − *x*) with the allowed domain *x < x*_0_ and for most problems *k* = 1. When *x << x*_0_ this function has negligible effect, however when *x* approaches *x*_0_ from below (inside the wall) the barrier function imposes an impregnable wall. Note that the reciprocal restraint function would also be subject to the “black hole” effect if it was not for a fundamental difference between minimization and MD – the existence of momentum at finite temperature. We provide a thought experiment here without formal mathematical proof, but we shall demonstrate that reciprocal restraint is effective in real-world applications and has worked in a variety of complex MD simulations in our drug discovery projects. In both examples presented here (protein conformational transitions and passive membrane permeability) *x* represents a complex CV that is a nonlinear function of the atomic coordinates (see 2). So, assuming that the current value of *x* is violating the CV restraint, it is outside the wall, the barrier function will pull it back toward the wall. Without momentum the system would never get inside the wall, though, instead it would fall into the exterior infinite energy well, and before reaching the singularity at *x*_0_ the MD simulation would crash with numerical overflow. However, at finite temperature the overall momentum of the system will carry it across the wall before falling into the singularity, and once inside, there is no escape. This means that we can start the simulation outside the wall, but it will quickly cross into the allowed domain and stay there indefinitely. Because the MD time step is finite, theoretically, the system can escape and end up outside, but even then it would quickly return as we suggested. Nonetheless, across a total of tens of microseconds of simulation time we have not experienced a single escape through our barrier wall.

## 2 Methods

We ran our simulations on Nvidia RTX 2080Ti and A40 GPU-equipped, Ubuntu 20.04 workstations, using the OpenMM-PLUMED software [11–14], versions 7.6.0 and 2.7.1, respectively. The barrier function was defined in PLUMED as a custom bias. One collective variable (CV) involved a path representing the putative transition between the open and closed conformations of the full CRL-VHL-degrader-SMARCA2 ring complex [15]. There are two common types of path CVs in use, one by Branduardi and Parrinello [16] (BP) and one by Leines and Ensing [17, 18] (LE). Both path variables are comprised of an ordered set of equally spaced intermediate conformations called nodes between two end conformations. Note, however, the node to node distance as well as the distance between the current simulation point in phase space and a path node can be quite different depending on what distance measure is used. Collective variables based on the root-mean-square-distance (RMSD) of the region of a protein known to be relevant to a particular conformational change, have been used [19, 20], but here our CV is the path and we use RMSD to compute the distance from the path nodes. We have experimented with both types of path variables extensively and we ended up using the BP path. When using the LE path we noticed that (1) the 3 closest path-nodes with respect to the current simulation point must be consecutive nodes, and (2) the closest node must be the middle node. In our simulations the second condition failed relatively frequently causing a discontinuity in computing how far along the path the simulation progressed, which led to the simulations crashing. As we discuss below, the BP path can exhibit similar discontinuities, but the barrier restraint described and implemented in this work is able to overcome that.

Equally spaced nodes were generated by an in-house minimum-energy path algorithm that extends Benoît Roux’s anisotropic network model (ANM) [21]. We are working on a separate publication focusing on the details of our path algorithm, but it is instructive to note here that the main difference as shown on Figure 1 is that our ANM model is smooth and differentiable everywhere, whereas the Roux path is based on a dual ANM model represented by two independent harmonic energy wells (one for each endpoint conformation) separated by a non-differentiable border where the two models intersect (see white border in the lower part of Figure 1. Our smooth-ANM path is based on a unified ANM model, which features a genuine saddle point connecting the two energy wells as shown in the upper part of Figure 1.

**Figure 1:**
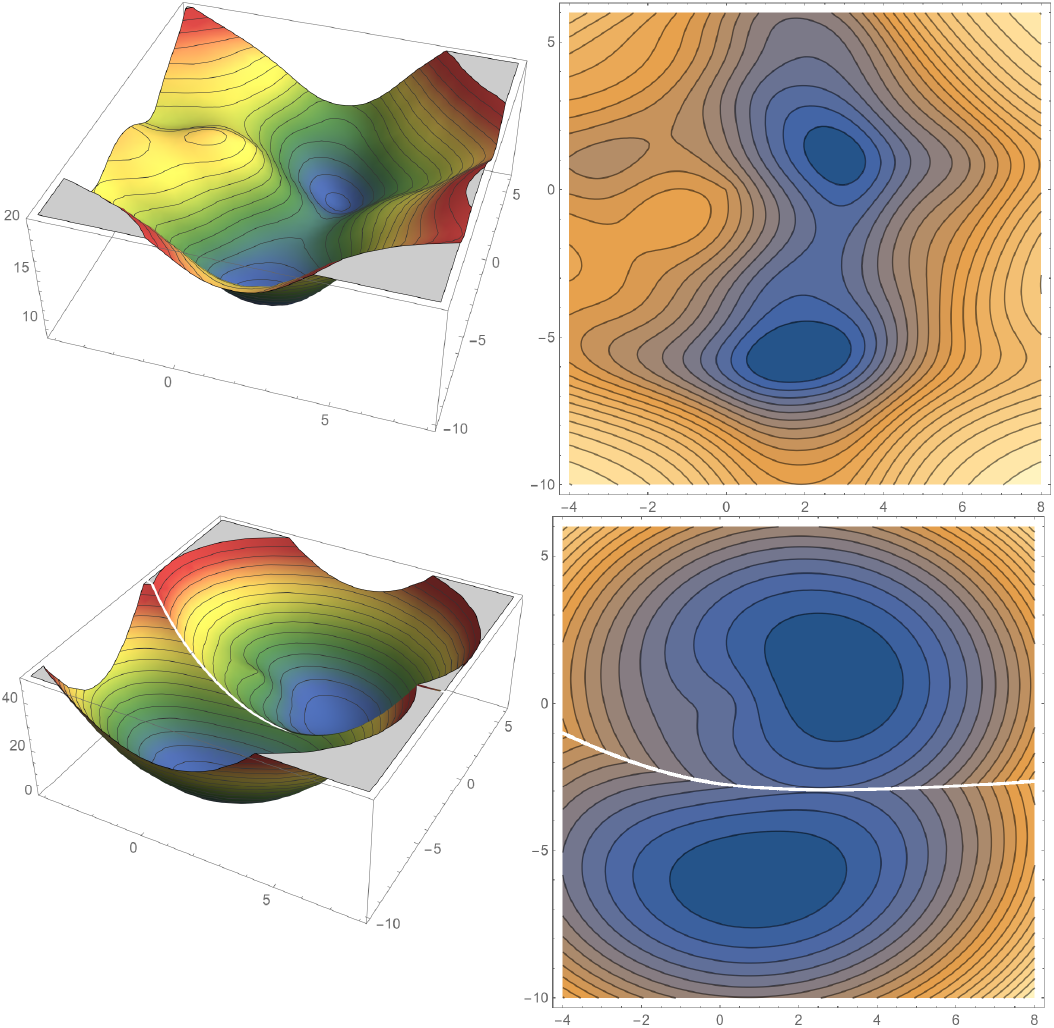
Smooth ANM minimum energy path The potential energy surface represents a hypothetical triangle shaped molecule with two distinct conformations corresponding to two different pairs of bond angles. The upper part shows our smooth ANM model with a genuine saddle point connecting the two energy wells, and the lower part shows the original ANM model [21] with its characteristic singularity, see text for details.

For biased simulations we employed the recently proposed combination of metadynamics and extended-Lagrangian adaptive biasing force (meta-eABF) method [22–24] for both biasing the simulation along the path and estimating the potential of mean force (PMF) of a large ligand molecule passing through a membrane model. Similar to ABF methods, meta-eABF simulations also utilize adaptive free energy biasing force to enhance sampling along one or more collective variables, but the practical implementation is different. Meta-eABF evokes the extended Lagrangian formalism of ABF whereby an auxiliary simulation is introduced with a small number of degrees of freedom equal to the number of CVs, and each real CV is associated with its so-called fictitious counterpart in the low-dimensional auxiliary simulation. The real CV is tethered to its fictitious CV via a stiff spring with a large force constant and the adaptive biasing force is equal to the running average of the negative of the spring force. The biasing force is only applied to the fictitious CV, which in turn “drags” the real simulation along the real CV via the spring by periodically injecting the instantaneous spring force back into the real simulation. Moreover, the main tenet of the meta-eABF method is employing metadynamics (MtD) or well-tempered metadynamics (WTM) to enhance sampling of the fictitious CV itself. The combined approach provides advantages over both MtD/WTM and eABF [25] alone.

In order to introduce our barrier restraint we first provide a closer look at the traditional approach. For sake of argument we look at an upper wall restraint located at 1. In other words, we do not want some variable *x* to go above 1. This upper wall is represented as a black vertical line in Figure 2. On the top right we show four different penalty functions using powers 2 to 5 (blue, orange, red, and green, respectively), and for simplicity we choose *F*_*c*_ = 1. Θ is the unit step function. The traditional quadratic penalty is represented by the blue curve. There are two features to note: (1) the penalty only kicks in for *x >* 1, and (2) these kind of penalty functions have very slow acceleration and therefore grow very slowly. In fact, the faster, higher powered functions start making a significant difference only for *x >* 2. Penalty functions such as this are only good for weak restraints in MD simulations. The negative derivative of the penalty function/potential is the restraining force. Figure 2 (bottom right) shows the forces associated with the penalty potentials. The forces are negative, pulling *x* back toward 1.

**Figure 2:**
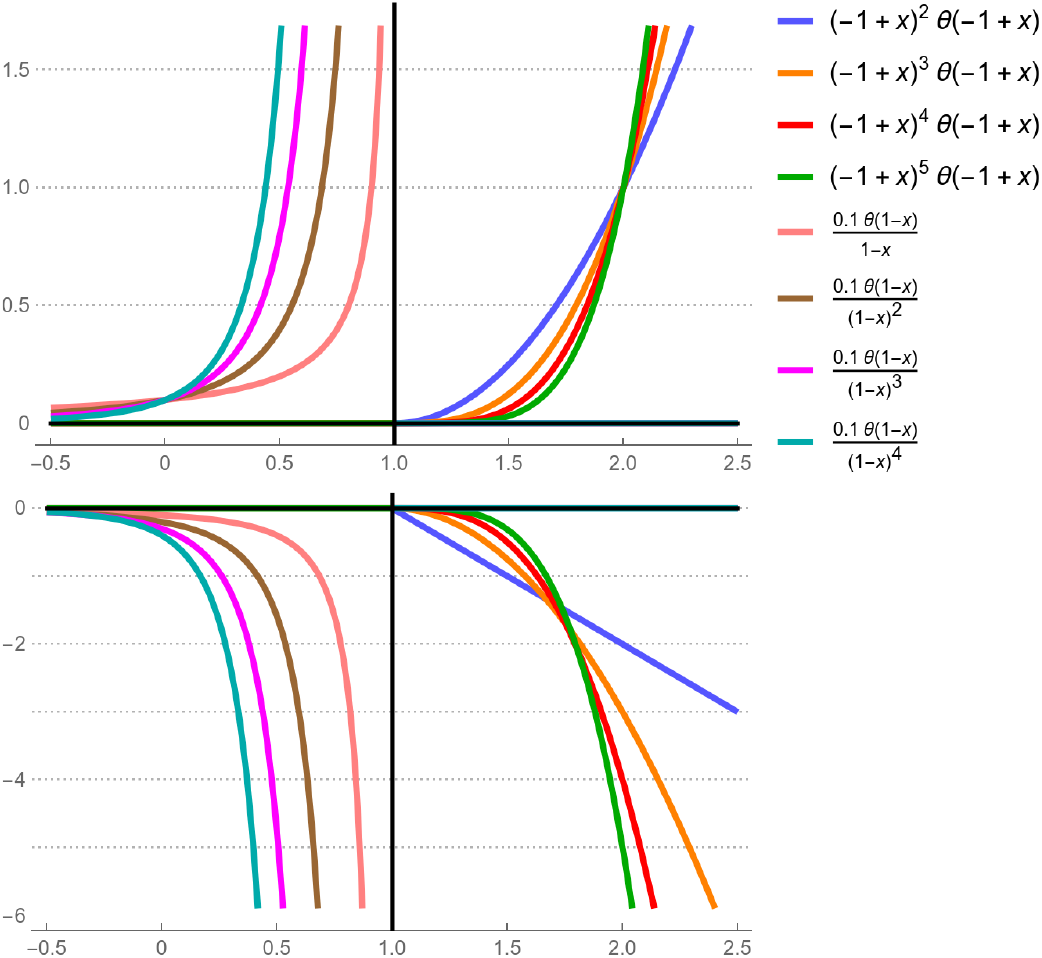
Barrier vs. penalty restraint The upper part shows the restraint potential of a series of barrier restraints (left) and penalties (right), and the lower part shows the corresponding restraint forces. The upper wall is represented by the vertical black line located at 1.0, see text for details.

From a practical perspective, it would be preferable for the penalty function to engage inside the wall and not outside. Barrier functions are designed to never let the restrained variable escape from its allowed domain by applying an infinite wall. Figure 2 (top left) shows a series of such barrier functions that are actually related to the penalty functions, but instead of using positive powers, they use negative powers. In general, we apply a simple and robust barrier function of the form 1*/*(*x*_0_ − *x*), which is used in all of the path simulations presented in this work, although for visual purposes the barrier functions are scaled by 0.1 in Figure 2.

Barrier functions are never zero in the allowed domain, but their values are very small up until they get close to the wall where they grow exponentially fast and go to infinity in the limit. Higher negative powers present a steeper wall and allow *x* to get closer to the wall. However, similar effect can be achieved with the lower powers by multiplying them with a small constant *k <<* 1, which we did in our membrane simulations. Figure 2 (bottom left) shows the barrier forces. Again, the forces are negative, pulling *x* away from the wall, keeping it inside. Also note how much larger the magnitudes of the barrier forces are in the vicinity of the wall with respect to the penalty forces on the bottom right. We shall give specific reasons why barrier restraints were superior to penalties in our detailed examples below.

All simulation job files for the work presented here have been submitted to the PLUMED-NEST repository [26]. For reference, the difference between penalty and barrier restraints in PLUMED syntax is shown below for an arbitrary upper and lower limit.

~~~
UPPER_WALLS ARG=my_x AT=3.0 KAPPA=250 LABEL=uwall_my_x
LOWER_WALLS ARG=my_x AT=1.0 KAPPA=250 LABEL=lwall_my_x
CUSTOM ARG=my_x FUNC=1/(3.0-x) PERIODIC=NO LABEL=up_x
BIASVALUE ARG=up_x LABEL=uwall_my_x
CUSTOM ARG=my_x FUNC=1/(x-1.0) PERIODIC=NO LABEL=lo_x
BIASVALUE ARG=lo_x LABEL=lwall_my_x
~~~

## 3 Results and Discussion

We recently published our work on atomic-resolution prediction of degrader-mediated ternary complex structures[15] where we also studied the dynamic nature of the full CRL-VHL-degrader-SMARCA2 complex via meta-eABF and generated a diverse set of putative closed conformations that place the E2-loaded ubiquitin close to lysine residues on SMARCA2. The results from the meta-eABF simulation are used to seed additional simulations for unbiased ensemble-scale sampling [15]. It was this meta-eABF path CV simulation where we first had encountered a serious problem with penalty restraints and found a solution using the barrier restraint described in this work.

Figure 3 shows the individual nodes representing discrete conformations along the path. The closed conformation is red, the fully open conformation is purple, and the stack of intermediate conformations are taking their colors from a rainbow palette. One can visualize the path CV as “morphing” between the two endpoints, but the minimum energy path is significantly more realistic than a geometric morph.[21]

**Figure 3:**
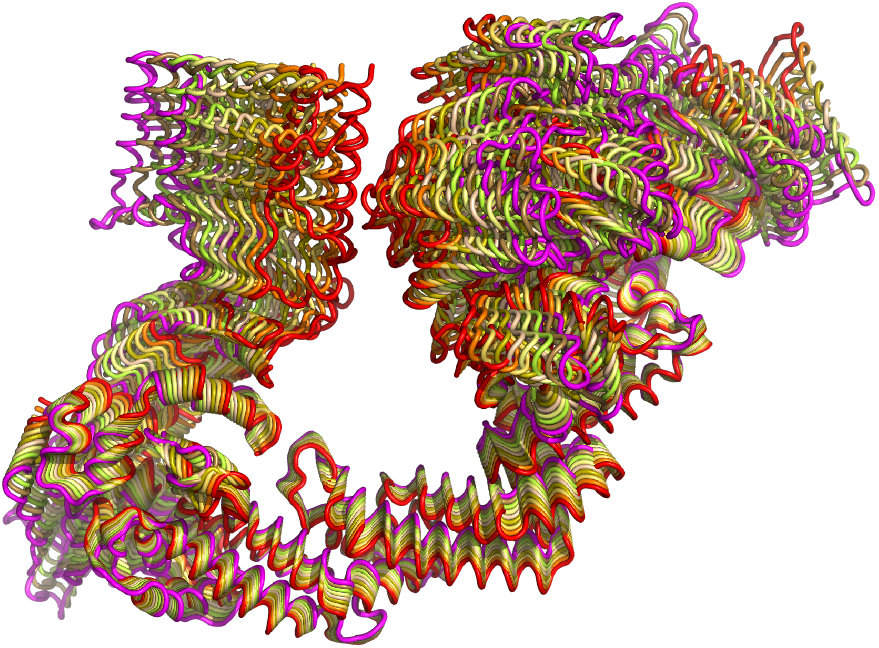
Frame stack representing the path CV of opening and closing the full CRL-VHL-degrader-SMARCA2 complex The closed conformation is red, the fully open conformation is purple, and the stack of intermediate conformations are taking their colors from a rainbow palette.

The computation of the path CV is quite complex [16] even though it is represented by only two degrees of freedom. One CV is the *S* variable representing the progress along the path so, e.g., if we have 50 nodes then *S* goes from 1 to 50. The other CV *Z* represents the orthogonal distance from the path. If we put an upper limit to *Z, S* and *Z* together can be visualized as a tube with length *S* and width *Z*. The essence of modeling the open and closed transition, then, is running a MD simulation such that by applying meta-eABF bias to the *S* collective variable the simulation is being nudged to follow the path back and forth but staying inside the tube, which is exactly where the new kind of restraint presented in this work shows its strength.

The BP path algorithm is strictly exact only for very thin tubes, but the authors state that “(we) expect that for most physically relevant trajectories, and assuming that the path consists of a sufficiently large number of nodes, the method works” [16] (FIG. 1.). In our experience with protein systems the radius of the tube should not exceed 2.5-3 Å. The exact limit depends on the system and the path, but this estimate has worked across a broad range of simulations and different systems. If the simulation deviates further from the path, the computation of *S* and *Z* becomes indeterminate causing discontinuities in the ABF forces and simulations will crash. To address this, we limit the tube radius to ∼3 Å (based on empirical observations). In our experience, applying a quadratic upper wall at 3 Å (i.e. the traditional approach) does not work, and even if we apply a large force constant, the simulation will make excursions beyond 3 from the path and will invariably crash. Even if we apply an offset and start the penalty at *Z* = 0 and/or use a *>* 2 power, there is no way of keeping the simulation inside the 3 Å tube using penalty restraint. If we tried increasing the force constant significantly to keep the system within the desired bounds then the large force constant itself caused the simulations to crash. As a rule of thumb, simulations usually cannot survive if the net force on a single atom exceeds 10^4^ kcal/mol/Å. In the actual meta-eABF simulation using the penalty restraint we applied the quadratic function centered at *Z* = 0 with *F*_*c*_ = 250 kcal/mol/Å^2^. Larger force constants crashed the simulation almost instantly, while with this value the simulation made excursions up to 4 Å from the path and crashed all the same.

Figure 4 shows the evolution of the *S* variable in a slice of the meta-eABF simulation of the CRL-VHL-degrader-SMARCA2 system from 30-60 ns. The vertical axis uses the serial numbers of the path CV nodes as units, but they roughly correspond to Angstroms, with values close to 1 representing the closed conformation and close to 40 representing the open conformation. First, consider the purple plot, which corresponds to using the traditional restraint on *Z*. It exhibits several spikes where *S* jumps several units, showing clear evidence of discontinuity. The simulation survives for a while but eventually crashes at about 55 ns. In Figure 5 we zoom in the middle to show the spikes more vividly.

**Figure 4:**
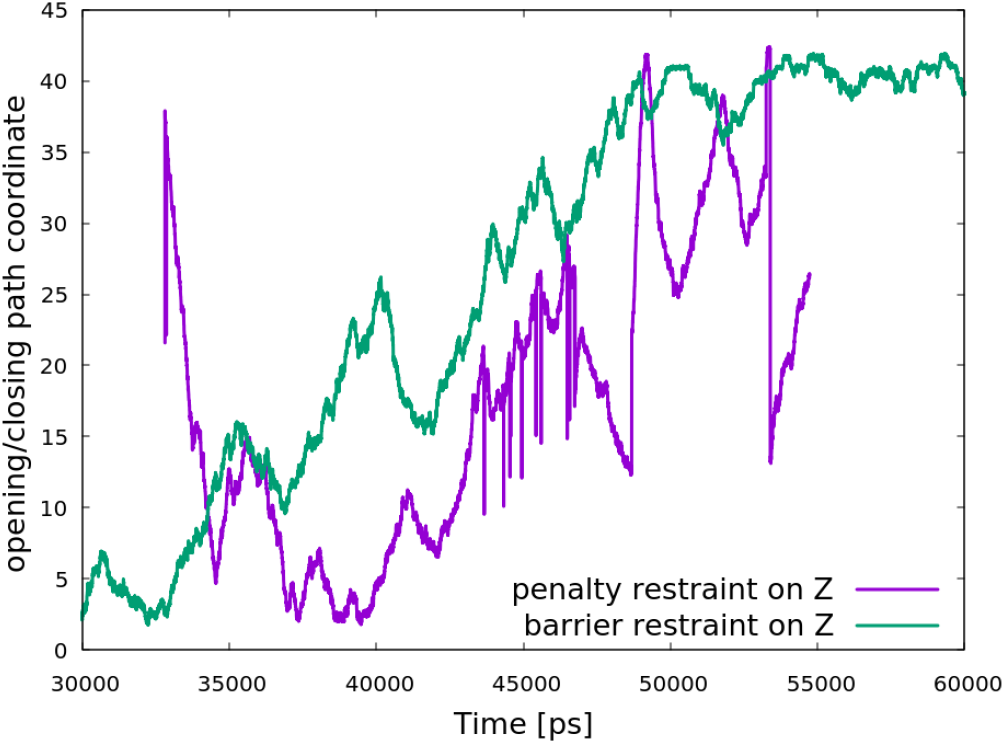
Time evolution of the *S* path variable Using harmonic penalty restraint on the *Z* variable (purple), the simulation has significant jumps along the path shown by the spikes along the vertical *S* coordinate, and crashes at about 55 ns. However, using our reciprocal barrier restraint (green), the simulation is smooth and runs indefinitely, see text for details.

**Figure 5:**
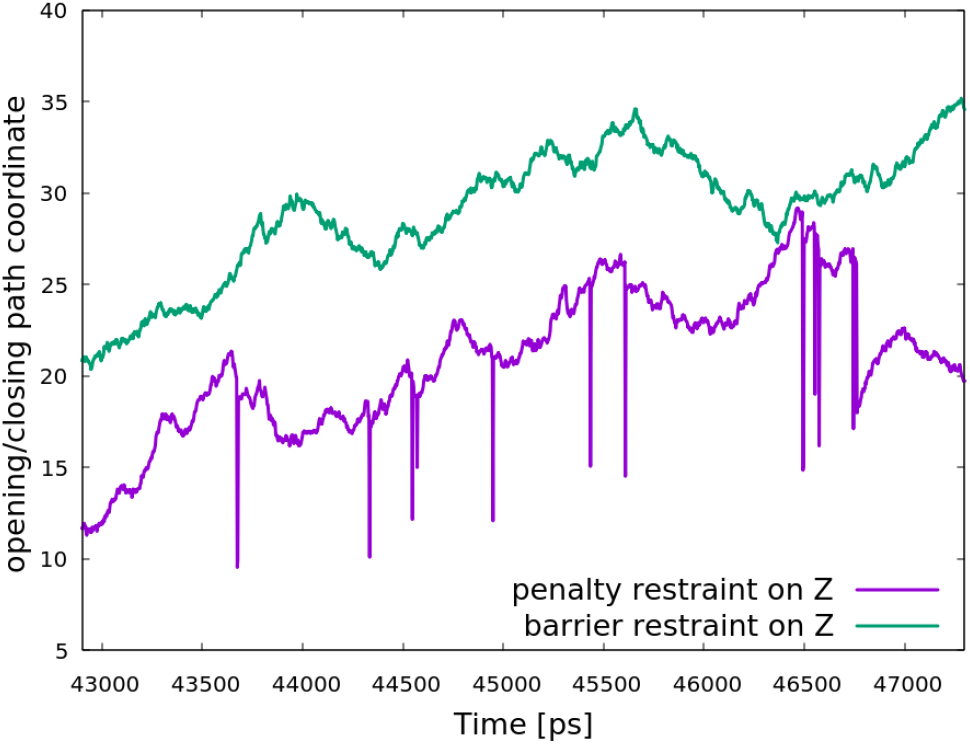
Time evolution of the *S* path variable, zoomed Same as Figure 4 but the time dimension is zoomed to a narrow slice to emphasize the jumps along the path when using penalty restraint for the tube radius *Z*.

Figures 6 and 7, respectively, show the same issue in the *Z* dimension. The purple spikes are nonsensical— as mentioned above, the *S* and *Z* spikes originate from a mathematical formula [16] that breaks down at about *Z* = 3 Å. As spectacular the spikes are, the important message in this figure is actually the approximate value of *Z* where there are no spikes, somewhere between 34 and 43 ns. It is hard to see it on Figure 6, but Figure 7 shows that the “normal” *Z* values hover between 3-4 Å, which is way beyond the safe tube radius.

**Figure 6:**
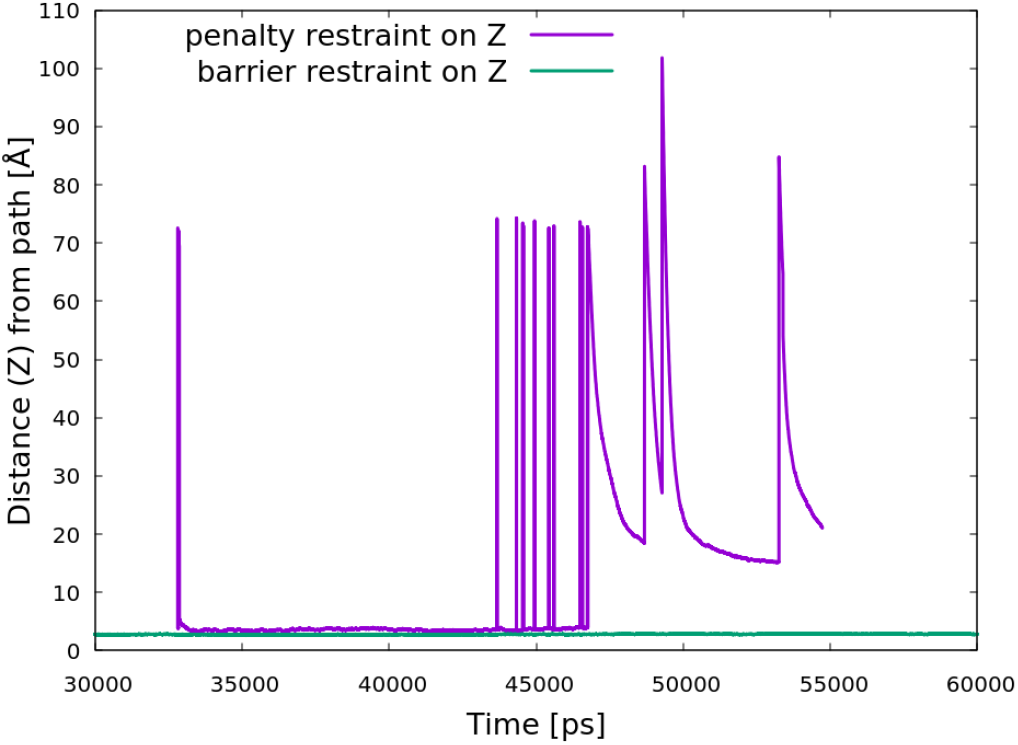
Time evolution of the *Z* path variable The jumps in the *S* variable shown in Figure 4 are coupled with enormous jumps in the *Z* variable. See text for details.

**Figure 7:**
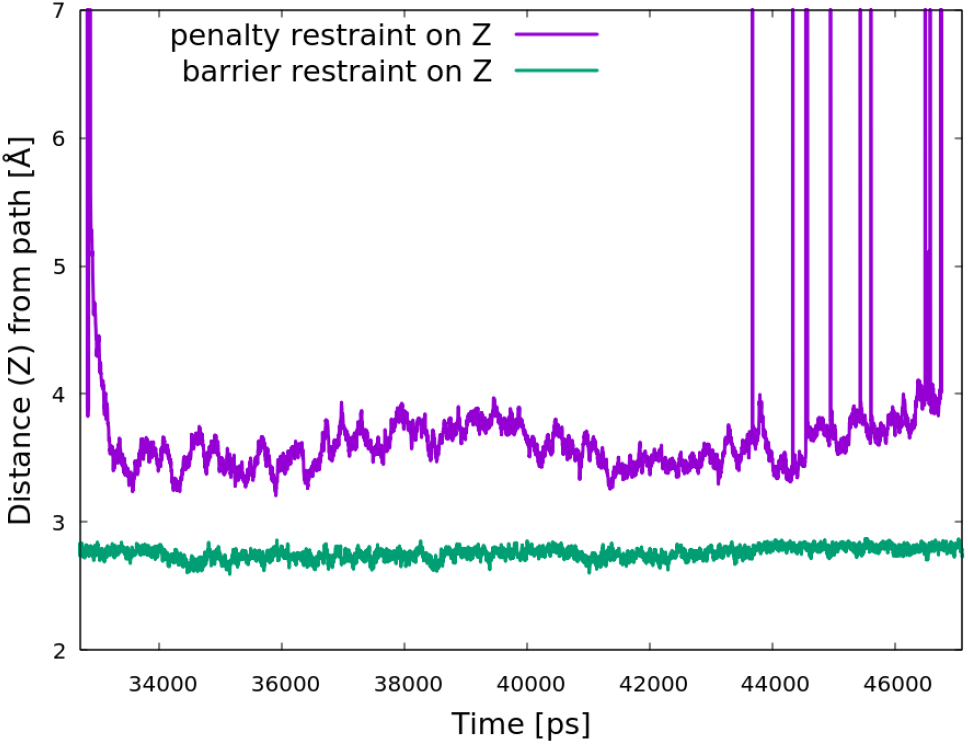
Time evolution of the *Z* path variable, zoomed After cropping the nonphysical *Z* range it becomes clear that even the very strong harmonic restraint that actually starts at *Z* = 0 (see text for details) cannot keep the simulation within the 3 Å tube whereas with our reciprocal barrier restraint *Z* stays well within the set limit.

Now, consider the green plots representing the exact same simulation except for using the barrier restraint instead of penalty. It is striking that both *S* and *Z* are perfectly smooth (the plots show data points at 1,000 MD time step intervals), *Z* stays below 3 Å and the simulation runs without incident for the set time of 100 ns.

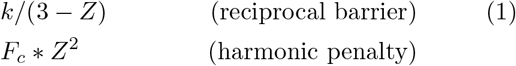

We achieved this by employing the reciprocal barrier function applied to the *Z* tube radius as shown in Equation 1 where *k* = 1 kcal/mol*Å and the tube is set to have a *Z* = 3 Å radius. In Figure 8 we compared the restraint potentials and forces for barrier vs. penalty. The horizontal axis represents the tube radius *Z*, shown only at the vicinity of the 3 Å limit where we want the restraint to behave like a hard wall. The positive domain of the vertical axis shows the restraint potential in kcal/mol and the negative domain represents the restraint force in kcal/mol/Å. The orange curve is the harmonic penalty potential with *F*_*c*_ = 250 kcal/mol/ Å^2^ with its value equal to 2,250 kcal/mol at *Z* = 3 Å. The harmonic potential has very little curvature, it is almost linear in this range and makes virtually no difference whether inside our outside the wall. Also note that the penalty starts at *Z* = 0 and therefore has a substantial value close to the wall, but as Figure 7 (purple curve) shows, the simulation still proceeds way outside the wall, up to 4 Å from the path. In stark contrast, the blue barrier restraint has negligible value all the way up to the immediate vicinity of the wall where it provides an infinite barrier preventing the simulation from ever leaving the tube. In fact, with *k* = 1 kcal/mol*Å the barrier restraint kicks in much earlier and as shown on Figure 7 (green curve), the simulation stays within ∼2.8 Å of the path. Note that by utilizing a small *k <<* 1, theoretically the simulation can get arbitrarily close to the wall, but there are numerical limits, which we shall discuss below with respect to our membrane simulations. Finally, the lower part of Figure 8 tells the same story showing the penalty force (green) and the barrier force (red), respectively. The restraint force is the manifestation of the wall—the barrier force represents a hard wall whereas the penalty force represents a soft wall.

**Figure 8:**
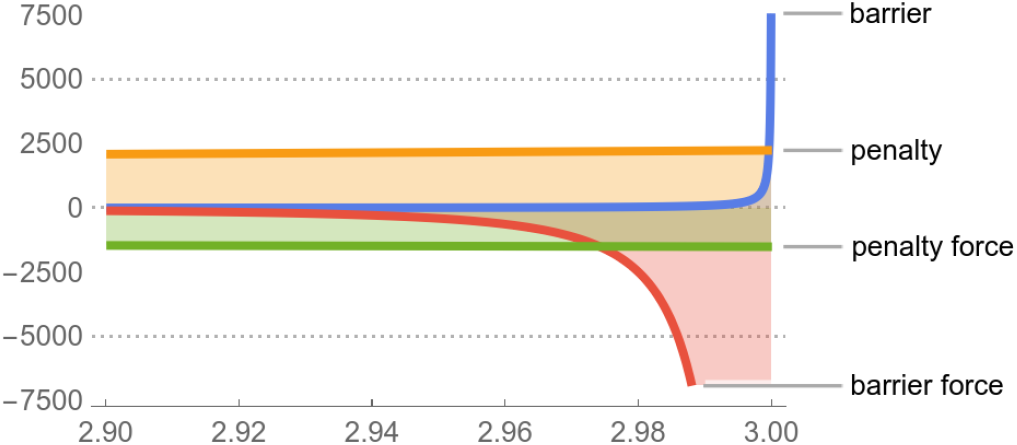
Path tube restraint: barrier vs. penalty Here we compare the harmonic penalty with our reciprocal barrier restraint. The horizontal axis represents the tube radius *Z*, shown near the 3 Å limit where we want the restraint to behave like a hard wall. The positive domain of the vertical axis shows the restraint potential in kcal/mol and the negative domain represents the restraint force in kcal/mol/Å. The orange curve is the harmonic penalty potential and has very little curvature, it is almost linear in this range and makes virtually no difference whether inside our outside the wall. The penalty starts at *Z* = 0 and therefore has a substantial value close to the wall, but as Figure 7 (purple curve) shows, the simulation still proceeds way outside the wall. In stark contrast, the blue barrier restraint has negligible value all the way up to the immediate vicinity of the wall where it provides an infinite barrier preventing the simulation from ever leaving the tube. See text for more details.

Heterobifunctional degrader molecules (also known as proteolysis targeting chimeras, or PROTACs) induce degradation of target proteins through the proteosome, which is present inside cells. As such, PROTAC molecules must have a sufficient level of cellular permeability to enter the cell. The bifunctional composition of PROTACs (two different protein binding ligands attached by a linker) creates challenges in obtaining desirable drug-like properties, where permeability is a critical factor. The large size of these molecules coupled with the relatively limited public data (the degradation field is relatively nascent) has limited the impact of computational approaches to predict permeability, although some learnings are starting to emerge [27, 28]. For this reason, we have focused on heterobifunctional degrader molecules for one case study in this work.

We have used meta-eABF to estimate the PMF for small molecule passive permeability in membrane models. An example of the results is shown in Figure 9. The CV in this case is the normalized and periodic *Z* coordinate of the center of mass (COM) of the propanol molecule with the domain [-0.5, 0.5] where the floating origin of the coordinate system corresponds to the COM of the membrane. Using a floating coordinate system has the advantage over a fixed origin that there is no need for tethering the membrane COM to an absolute point in the simulation box. Encouraged by our initial results, we employed the same protocol to much larger molecules ACBI1, PROTAC1, and PROTAC2 shown on Figure 10, but the results with the same protocol were poor (see Figure 11). The PMF curves are thermodynamically averaged from 4 independent, 1 *µ*s simulations for each molecule.

**Figure 9:**
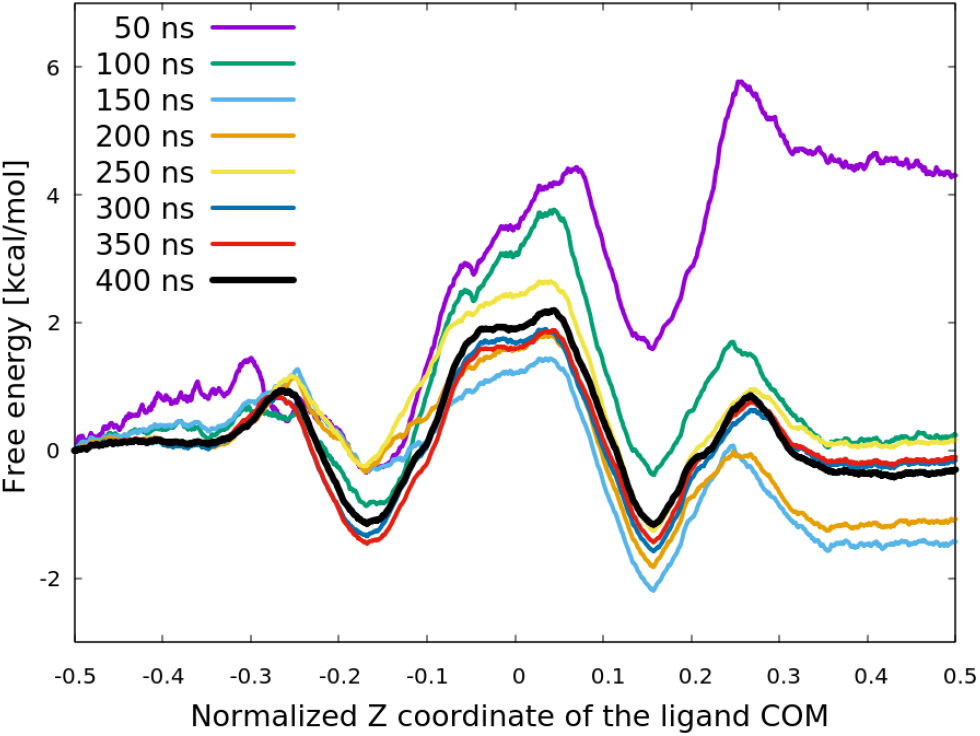
Time evolution of the estimated PMF of propanol permeation through POPC Initial 100 ns simulation is visibly not converged, but after 300 ns the PMF starts settling down, and by 400 ns it is converged showing the expected symmetric potential on either side of the center of the membrane.

**Figure 10:**
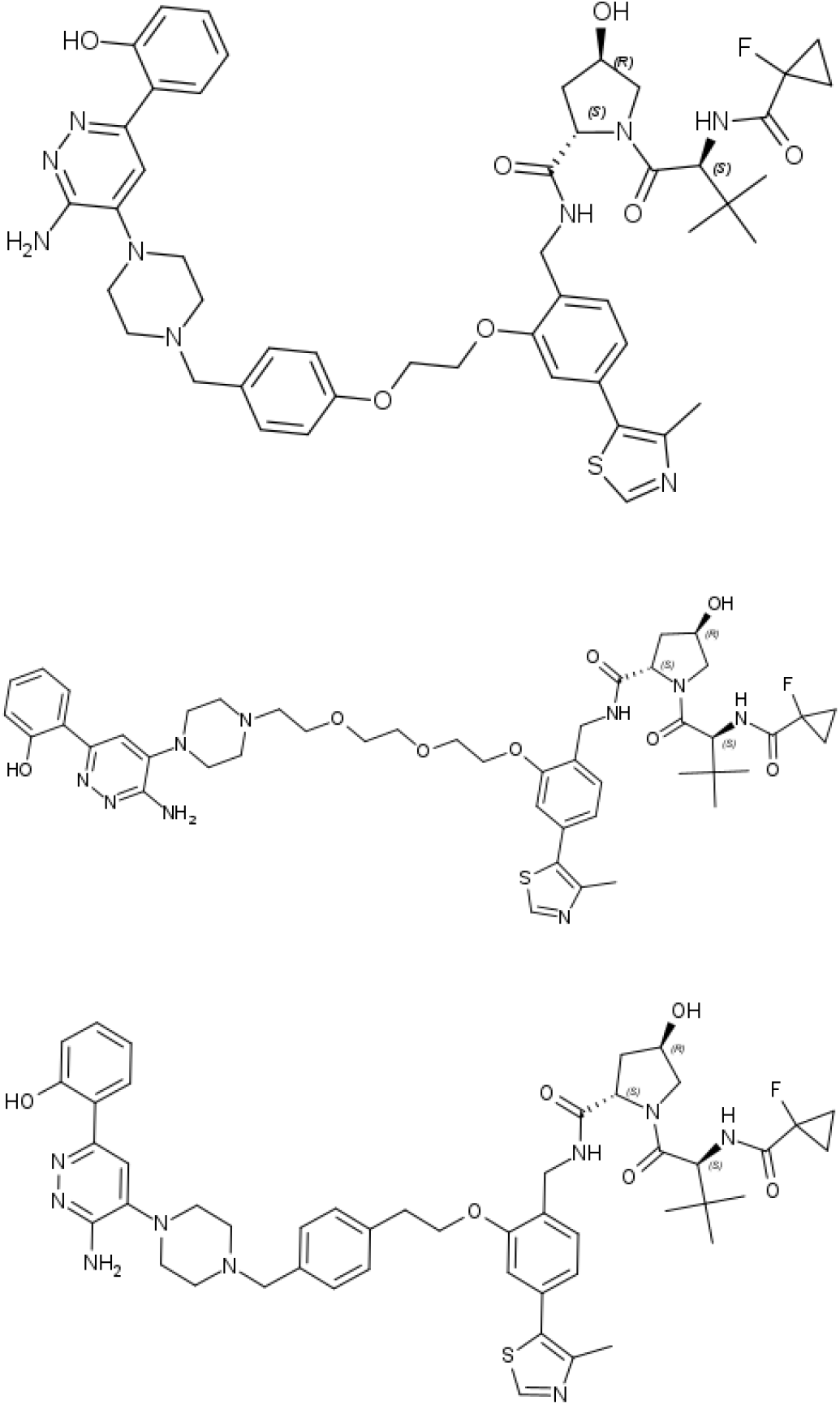
PROTAC ligands: ACBI1 (top), PROTAC1 (middle), PROTAC2 (bottom)

**Figure 11:**
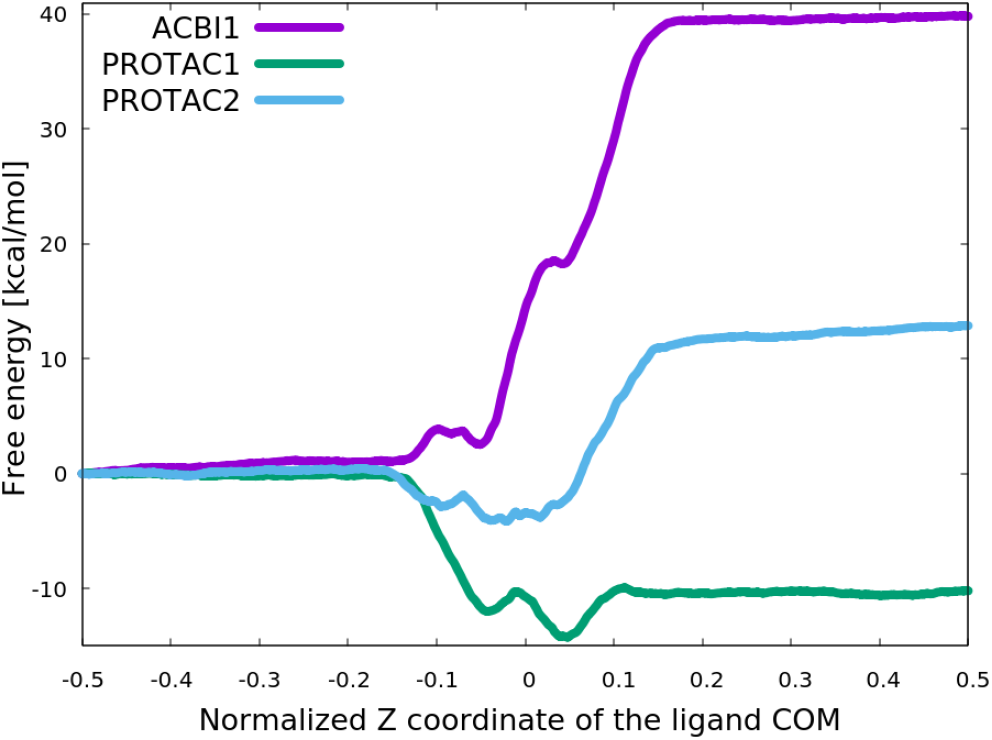
Naïve attempt to estimate the PMF of passive permeability of large heterobifunctional degrader molecules (ACBI1, PROTAC1, and PROTAC2) in a POPC membrane model Using the exact same protocol that provided acceptable results for propanol shown in Figure 9 turned out to be thoroughly inadequate for these large degrader molecules. The CV is the normalized and periodic *Z* coordinate of the center of mass (COM) of each of the degrader molecules with the domain [-0.5, 0.5] and the floating origin of the coordinate system corresponds to the COM of the POPC membrane. The PMF curves are thermodynamically averaged from 4 independent, 1 *µ*s simulations for each molecule, but the free energy estimates are very poor.

Visual inspection of the trajectories prompted our hypothesis that the spurious results were arising because these molecules spent too little time inside the membrane for adequate sampling across the entire permeation path. For example, propanol spent ∼ 60% of simulation time inside the membrane whereas ACBI1, PROTAC1/2 spent *<* 20%. For one, these large molecules carry a lot of momentum that helps propel them through the membrane quickly. While there are many ways to slow down this process (e.g. significantly delaying the meta-eABF bias), we could not find a satisfactory solution. To further encourage these large molecules to spend more time in the membrane, we found that the barrier restraint described here provided significant improvement.

To implement the barrier restraint in the membrane, we placed a double sided “mirror” in the middle of the membrane in the (*X, Y*) plane so that the COM of the ligand molecule bounces back from the mirror and does not let the molecule pass through the membrane from either up or down directions. In fact, given a perfect mirror, we only need one half of the simulation box because of the symmetry of the system in the *Z* dimension, but using a double sided mirror and the entire simulation box with periodic boundary conditions (PBC), allows for two simulations in one. One simulation is progressing under the mirror and another simulation above the mirror, and the system switches between the two simulations via PBC through the water phase. Meta-eABF data collection is disconnected in the “up” and “down” simulations using two independent CVs, one is the *Z* coordinate of the ligand COM in the domain [-0.5, 0) and the other (0, 0.5]. We do not have a perfect mirror, but the barrier restraint comes close, shown in Equation 2. It should be noted that the free energy associated with orientational rearrangements at the center of the membrane are not captured in this framework, which is a known limitation and something we will explore in future works.

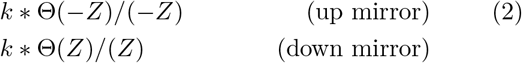

There are two important characteristics of the mirror function, one is that *k* should be set to as small as numerically possible to mimic a vertical wall. With *k <<* 1 there is virtually no restraint inside the wall and we can get arbitrarily close to the wall. In our experience using single precision GPU code, *k* ≈ 0.008 is the smallest value that does not cause numerical artifacts, such as the COM of the ligand molecule sticking to the mirror. The other characteristics is the requirement of including the Θ unit step function. We explained above that reciprocal barrier functions do not inevitably break down in MD simulations upon leaving the interior domain, and that is quite beneficial using upper and lower walls that are separated by a finite distance, i.e. a flat-bottom restraint, but in case of our double sided mirror the upper wall (up mirror, reflecting upward movement) and lower wall (down mirror, reflecting downward movement) coincide, and we do not want restraint forces to spill over to the other side.

Figure 12 explains the structure of the double sided mirror with respect to Equation 2 and *k* = 0.008. The horizontal axis is the normalized and periodic *Z* coordinate of the ligand COM between [-0.5, 0.5] and zero representing the floating middle point of the membrane. The membrane stretches between ∼ [-0.12, 0.12] and it is symbolically represented by the grey rectangle. The double sided mirror resides at *Z* = 0 and it is represented by the vertical line separating the colored regions. The upward movement of the ligand permeating through the membrane is left to right and downward movement is from right to left. The upper part of Figure 12 shows the restraint potential [kcal/mol] and the lower part shows the restraint force [kcal/mol/Å]. The potential and force are represented by the colored curves, the shading is just for visual emphasis. The colored horizontal line segments with zero potential/force value represent the zero branch of the unit step function.

**Figure 12:**
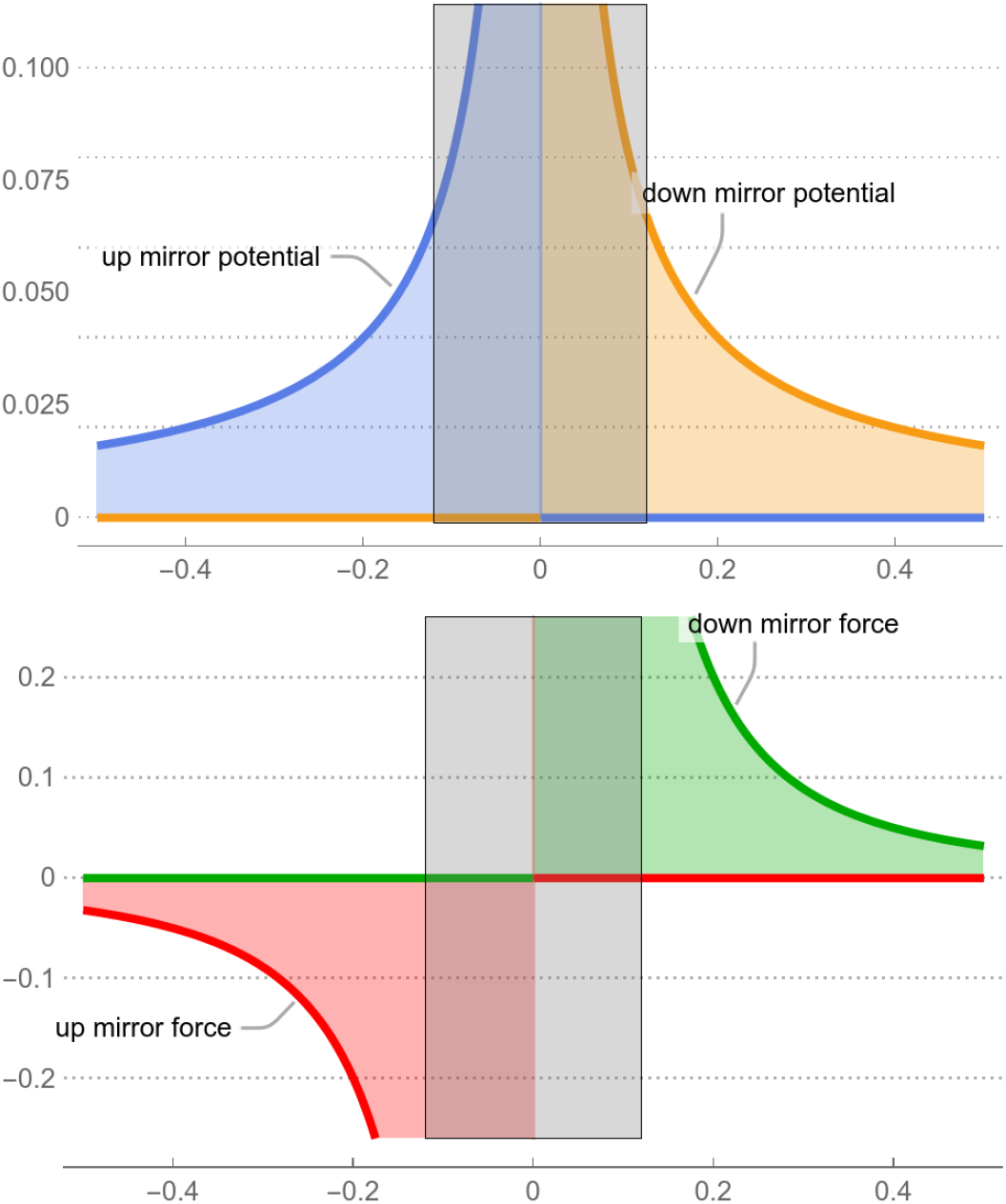
Double sided mirror restraint The double sided “mirror” restraint (Equation 2 with *k* = 0.008) prevents the ligand molecule from passing through the middle plane of the membrane from either side. The horizontal axis is the normalized and periodic *Z* coordinate of the ligand COM between [-0.5, 0.5] and zero representing the floating middle point of the membrane. The membrane stretches between ∼ [-0.12, 0.12] and it is symbolically represented by the grey rectangle. The double sided mirror resides at *Z* = 0 and it is represented by the vertical line separating the colored regions. The upward movement of the ligand permeating through the membrane is left to right and downward movement is from right to left. The upper part shows the restraint potential [kcal/mol] and the lower part shows the restraint force [kcal/mol/Å]. The potential and force are represented by the colored curves, the shading is just for visual emphasis. The colored horizontal line segments with zero potential/force value represent the zero branch of the unit step function.

Figure 13 shows a significant improvement in the PMF curves with respect to Figure 11 as a result of application of the double sided mirror manifest in Equation 2 and Figure 12. These were also quadruplets of 1 *µ*s simulations for each ligand molecule and their COM was inside the membrane ∼ 40% of the time. We already mentioned that bouncing off the mirror introduced new, e.g., rotational degrees of freedom and single precision did not allow *k <* 0.008 in Equation 2, so further theoretical as well as computational studies are needed. For example, the effect of the latter can be seen in Figure 13 where the tips of the PMF curves are somewhat ill shaped, because there was not enough sampling in the extreme vicinity of the wall. Double precision would allow for using *k <<* 1 and that would produce much better sampling extremely close to the mirror. This is work in progress, nonetheless, using the double sided mirror made such spectacular improvement that we strongly feel it has its own merit. In fact, the barrier height of the PMF curves (measured between the minimum and the maximum, inside the membrane) are in good qualitative agreement with experimental Caco-2 permeability measurements [29]: PROTAC1 has very low permeability, and ACBI1 and PROTAC2 have low/moderate permeability.

**Figure 13:**
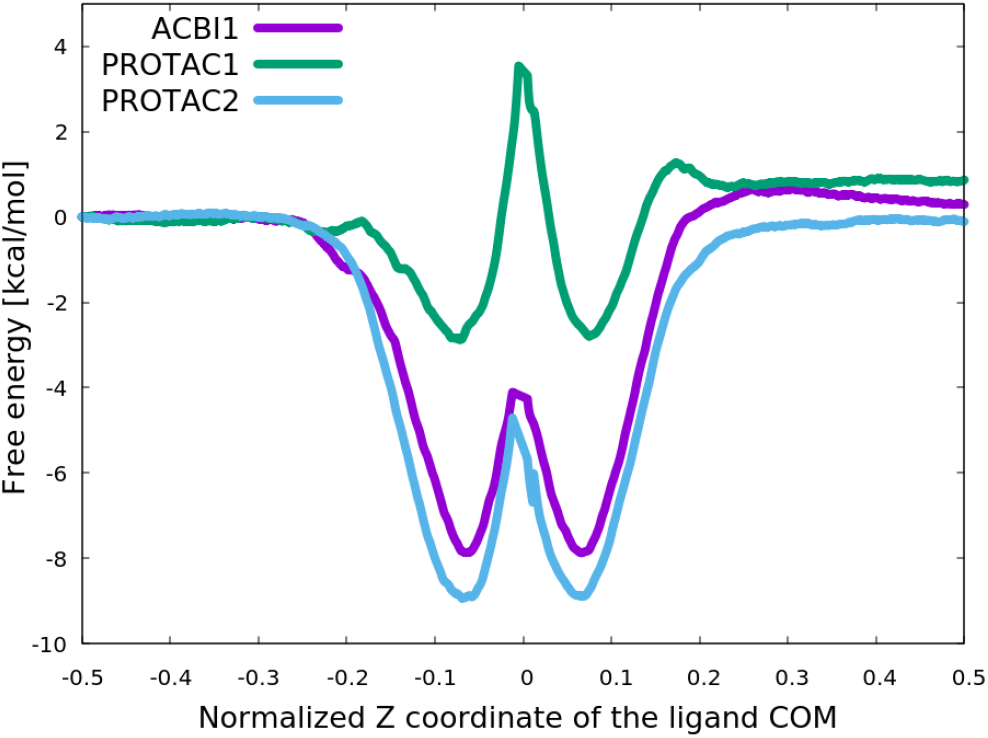
Improved estimates of the PMF of passive permeability of large heterobifunctional degrader molecules (ACBI1, PROTAC1, and PROTAC2) in a POPC membrane model Adding the mirror restraint (Equation 2 and Figure 12) to the protocol used in our naïve PMF estimates shown in Figure 11 resulted in compelling improvement. These were also quadruplets of 1 *µ*s simulations for each ligand molecule and the barrier height of the PMF curves (measured between the minimum and the maximum, inside the membrane) are in good qualitative agreement with experimental Caco-2 permeability measurements [29]. See text for more details.

Finally, we provide the entire PLUMED input below for running similar simulations. The list of atoms included in the COM calculations is stored as strings in `lig_com_heavy_atoms_str and mem_com_atoms_str`, respectively. Also note that the delayed meta-eABF bias is set by the `FULLSAMPLES=200000` parameter [30].

~~~
“““
UNITS LENGTH=A TIME=ps ENERGY=kcal/mol
COM ATOMS={0} LABEL=COM_lig
COM ATOMS={1} PHASES LABEL=COM_mem
DISTANCE ATOMS=COM_mem,COM_lig SCALED_COMPONENTS
  LABEL=COM_lig_dist
DRR ARG=COM_lig_dist.c FULLSAMPLES=200000 GRID_MIN=-0.5
  GRID_MAX=0 GRID_SPACING=0.001 TEMP=310 FRICTION=8.0
  TAU=0.5 TEXTOUTPUT OUTPUTFREQ=2500000 HISTORYFREQ=25000000
  DRR_RFILE=drr_lo LABEL=drr_lo
DRR ARG=COM_lig_dist.c FULLSAMPLES=200000 GRID_MIN=0
  GRID_MAX=+0.5 GRID_SPACING=0.001 TEMP=310 FRICTION=8.0
  TAU=0.5 TEXTOUTPUT OUTPUTFREQ=2500000 HISTORYFREQ=25000000
  DRR_RFILE=drr_up LABEL=drr_up
METAD ARG=drr_lo.COM_lig_dist.c_fict SIGMA=0.005 HEIGHT=1.5
  PACE=500 GRID_MIN=-0.5 GRID_MAX=+0.5 GRID_SPACING=0.001
  BIASFACTOR=8 TEMP=310 FILE=HILLS_lo LABEL=metad_lo
METAD ARG=drr_up.COM_lig_dist.c_fict SIGMA=0.005 HEIGHT=1.5
  PACE=500 GRID_MIN=-0.5 GRID_MAX=+0.5 GRID_SPACING=0.001
  BIASFACTOR=8 TEMP=310 FILE=HILLS_up LABEL=metad_up
CUSTOM ARG=COM_lig_dist.c FUNC=8e-3*step(-x)/(-x)
  PERIODIC=NO LABEL=mirror_up
BIASVALUE ARG=mirror_up LABEL=uw_mirror
CUSTOM ARG=COM_lig_dist.c FUNC=8e-3*step(x)/x
  PERIODIC=NO LABEL=mirror_lo
BIASVALUE ARG=mirror_lo LABEL=lw_mirror
PRINT ARG=* STRIDE=1000 FILE=plumed-meta-eabf-bias.log
FLUSH STRIDE=1000
“““.format(lig_com_heavy_atoms_str,mem_com_atoms_str)
~~~

## 4 Conclusion

In this work we presented a new kind of restraint for molecular dynamics simulations. While this restraint (we term it ReBaCon for Reciprocal Barrier Constraint) has significant challenges in nonlinear optimization, it is well-suited for molecular dynamics simulations. In particular, we have found it to be the best approach for path-based enhanced sampling simulations of complex biological conformational changes. In this work we have applied ReBaCon within a hybrid MD sampling method that combines metadynamics with extended adaptive biasing force (Meta-eABF). The method performs well for both membrane permeability of complex heterobifunctional degrader molecules and for large-scale conformational changes in the Cullin RING ligase macromolecular structure responsible for ubiquitination.

Accurate and efficient prediction of conformational changes (both the structures and the underlying free energy surfaces) is an important area of research with vast applications. While great strides have been made in protein structure prediction (e.g. AlphaFold, RoseTTAFold, and OpenFold), progress on conformational sampling has not witnessed a comparable acceleration from artificial intelligence (AI) and machine learning (ML), although some encouraging approaches are beginning to arise [31– 33]. One reason is that there is much less data associated with protein motion as compared with static structures. One of the few experimental methods that can assess protein motion and energies is NMR [34], which is challenging, time consuming, and limited data is available to date. We expect AI and ML to play an increasingly important role in conformational sampling, we suspect there will be a role for physics based simulations for the foreseeable future. Indeed, if physics-based simulations are accurate and efficient enough, they could provide the scale and quality of data necessary for an AI/ML approach.

In the approach presented here we have made an incremental improvement to a field rich with methods and applications—we are not proposing a step change in performance. Instead, we have chosen a very specific aspect of conformational free energy simulations (specifically, the restraining potential) and developed a solution that overcomes significant limitations of existing methods. We hope that ReBaCon will be used by others in the field to validate and improve the method, especially with other sampling methods and different biological systems. The implementation of this approach is simple and should facilitate quick testing and possible extensions. We encourage others in the field to try this method and propose further improvements, with a focus on biologically relevant problems as opposed to model systems (e.g. alanine dipeptide or Trp-cage). We are currently testing ReBaCon on a variety of conformational sampling problems, including protein-ligand binding, conformational activation, and allosteric inhibition.

## 5 Acknowledgements

IK is indebted to Siyoung Kim for discussions on simulating passive permeability, and Asghar Razavi and Wenchang Zhou for further discussions and help with system preparation. We also thank Jesus Izaguirre and Huafeng Xu for critical reading of the manuscript.

